# Integrative Ranking Of Enhancer Networks Facilitates The Discovery Of Epigenetic Markers In Cancer

**DOI:** 10.1101/2020.11.25.397844

**Authors:** Qi Wang, Yonghe Wu, Tim Vorberg, Roland Eils, Carl Herrmann

## Abstract

Regulation of gene expression through multiple epigenetic components is a highly combinatorial process. Alterations in any of these layers, as is commonly found in cancer diseases, can lead to a cascade of downstream effects on tumor suppressor or oncogenes. Hence, deciphering the effects of epigenetic alterations on regulatory elements requires innovative computational approaches that can benefit from the huge amounts of epigenomic datasets that are available from multiple consortia, such as Roadmap or BluePrint. We developed a software tool named Irene (Integrative Ranking of Epigenetic Network of Enhancers), which performs quantitative analyses on differential epigenetic modifications through an integrated, network-based approach. The method takes into account the additive effect of alterations on multiple regulatory elements of a gene. Applying this tool to well-characterized test cases, it successfully found many known cancer genes from publicly available cancer epigenome datasets.

## Introduction

Epigenetic alterations are frequent in many cancers. In particular, DNA methylation and histone modifications are two main mechanisms that allow cancer cells to alter transcription without changing the DNA sequences, and lead to many abnormalities such as persistent activation of cell cycle control genes or deactivation of DNA repair genes. For example, promoter DNA hypo-methylation accompanied by histone hyper-acetylation is frequently observed in the activation of oncogenes in cancer. Besides, aberrant activation of distal regulatory elements is often associated with the up-regulation of cancer-promoting genes. Interestingly, epigenetic modifications at proximal and distal regulatory elements often appear to be earlier events than the gene expression (Ziller et al., 2014; Hartley et al., 2013), and can hence serve as potential early markers in cancer diagnosis.

Various histone modifications on promoters have been categorized into either activation or repression effects on gene expression. Such effects can be measured by comparing histone alteration levels between tumor and their corresponding normal tissues using ChIP-Seq (Karlic et al., 2010). A number of tools, such as ChIPComp (Chen et al., 2015), ChIPDiff (Xu et al., 2008), ChIPnorm (Nair et al., 2012), csaw (Lun and Smyth, 2015), DBChIP (Liang and Keles, 2012), DiffBind (Stark and Brown, 2011), MAnorm (Shao et al., 2012), RSEG (Song and Smith, 2011) have demonstrated their usefulness in cancer studies by comparing the histone intensities between two conditions (see (Steinhauser et al., 2016) for a review of these tools). However, they are limited to the comparison of a single histone mark. Combinatorial effects of multiple histone marks are mainly performed in qualitative measurement. To the best of our knowledge, a method called differential principal component analysis (dPCA) is the only approach that performs quantitative analysis of multiple histone marks so far (Ji et al., 2013). It decomposes differential epigenetic signals between two biological groups from multiple histone marks and summarizes the variances from the largest to the smallest to several differential principal components (dPCs).

However, many histone modifications that potentially regulate gene expression also occur in the other genomic regions besides promoters. Enhancers are distal regulatory elements that interact with gene promoters through chromosomal loops to regulate gene transcription. Most of the enhancers are located within ±1 Mb of the transcription start site (TSS) of their target genes (Maston et al., 2006). Enhancer activity is regulated through epigenetic modifications(Zentner et al., 2011), including positive regulation from histone marks, such as H3K27ac (Creyghton et al., 2010; Stasevich et al., 2014) and H3K4me1 (Heintzman et al., 2007; Calo and Wysocka, 2013), and negative regulation by H3K27me3 (Charlet et al., 2016) and H3K9me3 (Zhu et al., 2012).

Given the complexity of epigenetic regulation, novel tools are required to combine this information, and create a comprehensive overview of the differential epigenetic landscape, integrating multiple data layers. The method we developed, named Irene (Integrative ranking with an epigenetic network of enhancers) combines a quantitative analysis on multiple differential epigenetic modifications with an integrated, network-based approach, in which we integrated two levels of epigenetic information: the signal intensity of each epigenetic mark, and the relationships between promoters and distal regulatory elements known as enhancers (Fig. 1). In this paper, we describe the method and present the test cases. In our benchmarking tests on cancer datasets, the Irene rank lists have higher relevance to cancer marker genes than the other approaches. Being implemented as an R package, Irene is an easy to use method allowing gene ranking between two conditions and highlighting potential cancer biomarkers.

**Figure 1:**
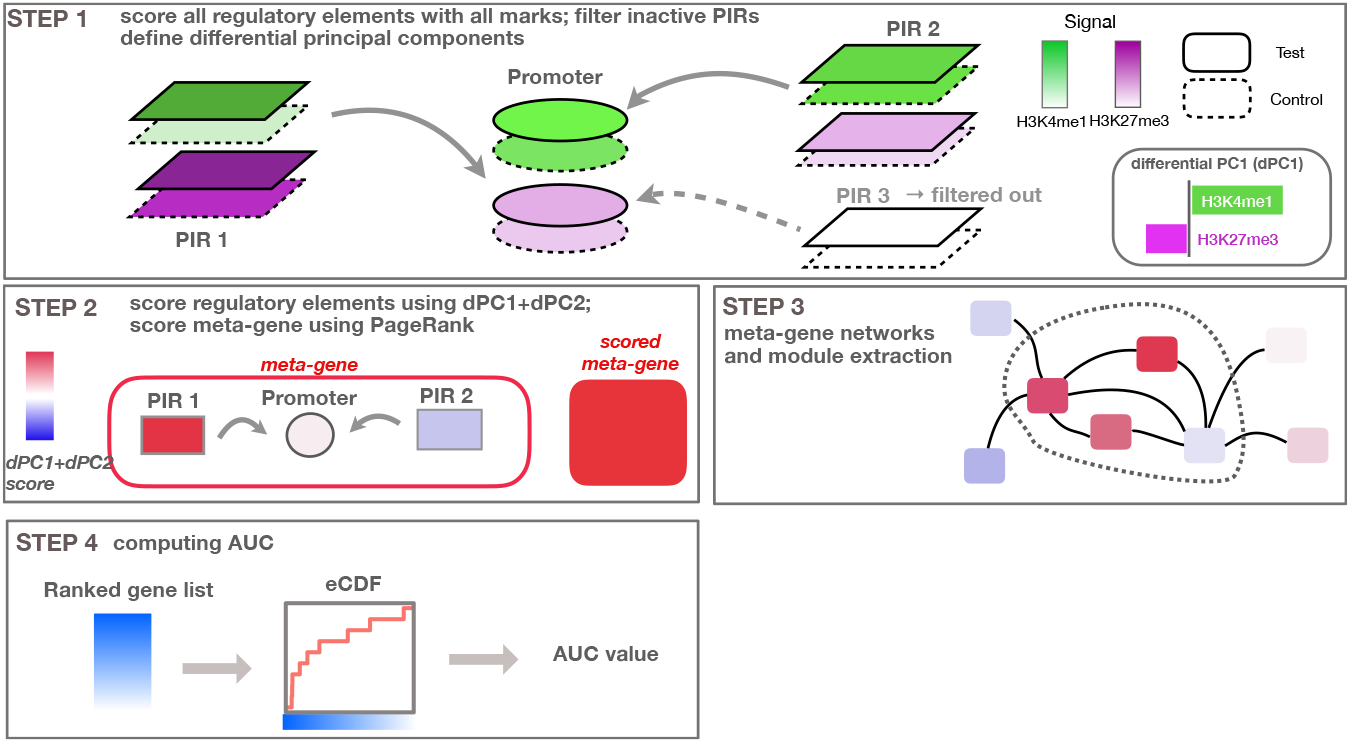
Overview of the method. (Step 1) Regulatory elements are scored using all epigenetic modifications, and related to the target gene. (Step 2) Epigenetic alterations are scored using the first differential principal component and combined using PageRank into an integrated meta-gene score. (Step 3) Functional interaction networks between meta-genes, based on protein-protein interaction data are used to extract sub-modules. (Step 4) Ranked gene lists based on the score are converted to eCDF curves showing the enrichment of a given gene set within the top-ranked genes, and corresponding AUC values are computed.

## Results

### Irene: Epigenetic ranking with an epigenetic network of enhancers

Irene analyzes epigenetic changes between two biological conditions (e.g. ChIP-seq data for histone modifications or whole-genome bisulfite sequencing for DNA methylation), and translates the differential signals at multiple regulatory elements into a unique score (Fig. 1). To integrate multiple datasets, we use differential principal component analysis (dPCA), which captures the directions of the greatest differential variance comparing two conditions, at each regulatory element. We consider both gene promoters as well as promoter interacting regions (PIRs) from the 4DGenome database. These scores are summarized as a weighted network relating regulatory elements to their target genes. A Random walk based method then assigns a score to the corresponding gene. The output of the method is a ranked list of genes from the most to the least affected one, which incorporates both promoter and enhancer alterations. This approach can be applied whenever two conditions are to be compared, for example, normal/tumor tissue, various tumor subtypes or different developmental stages. More details are given in the methods section. In order to benchmark our method, we used seven test cases consisting of tumor samples for seven different tumor types, and normal matching samples. The test cases comprise samples from acute myeloid leukemia (ALL), IGH-mutated/unmutated chronic lymphocytic leukemia (mCLL/CLL), colorectal cancer (CRC), Glioma, Multiple myeloma (MM), papillary thyroid carcinomas (PTC). The files used for the comparison can be found in Supplementary Table 1. For each of these test cases, we compiled a list of cancer marker genes (CMGs, Supplementary Table 2) from the literature, and considered housekeeping genes (HKGs) as controls.

### Cancer marker genes are scored higher by incorporating enhancer in the ranking

In our analysis, we determined that taking into account the first two dPCs is able to capture most of the differential variance for both activating and repressive epigenetic modifications (Fig. 2a,b). After comparing the dPC1+dPC2 values between the CMGs and HKGs in each test case, we found that the scores from CMGs are generally higher than the scores of the HKGs, both for enhancers as well as promoters. (Fig. 2c).

**Figure 2:**
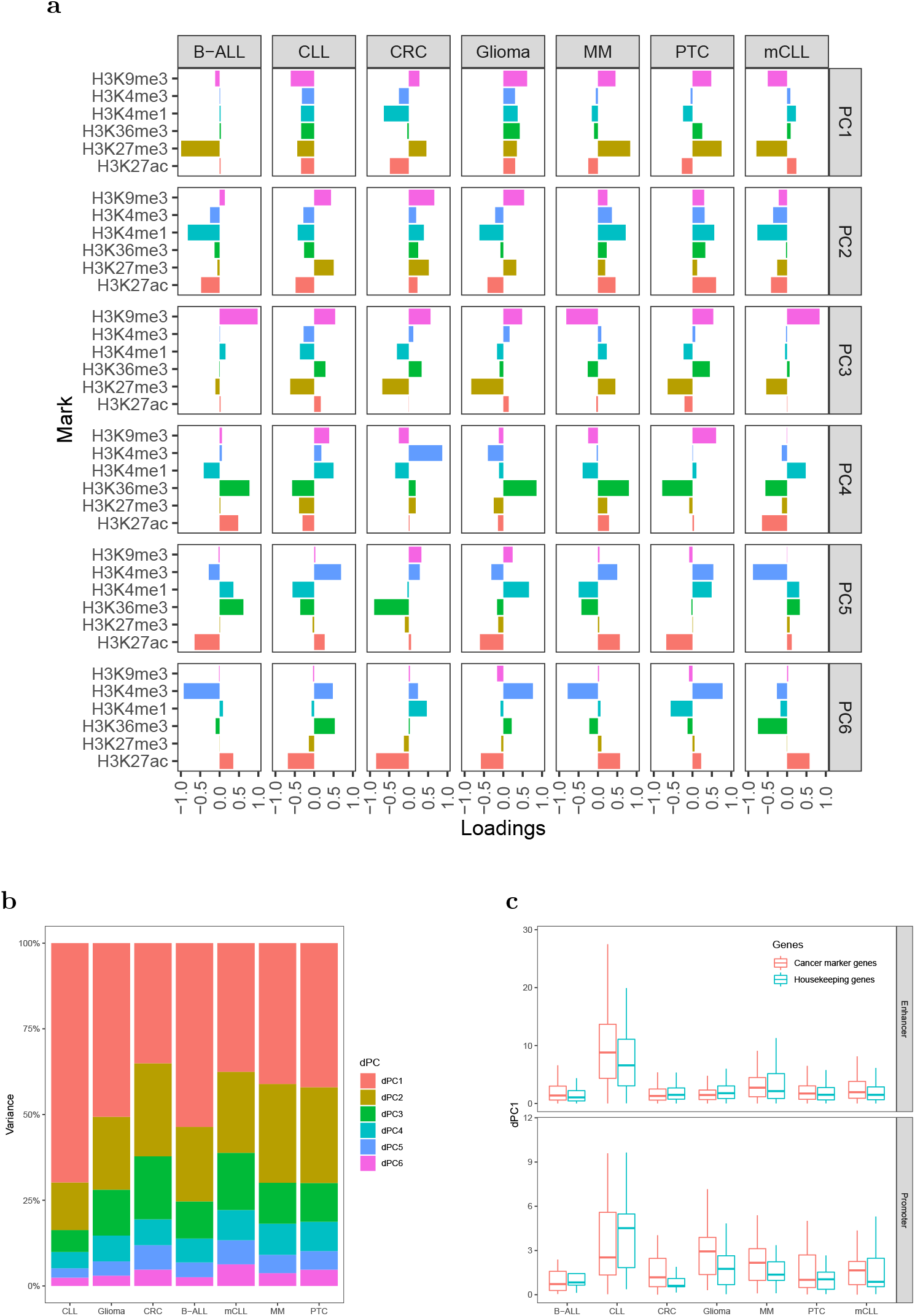
Differential principal components. (a) Contributions of the six histone marks to each dPC. (b) Variances accounted for each dPC in the seven test cases. (c) Values of dPC1+dPC2 in the seven test cases, comparing CMGs with HKGs, both for enhancers (top) and promoters (bottom).

We further computed the area under the curve (AUC) for the empirical cumulative density function (ECDF) of the high-confidence CMG ranks as a benchmarking approach, as described in the methods. First, we examined the Irene ranks computed using the dPC1+dPC2 on gene promoters and their targeting enhancers, and found that the marker genes are ranked higher than HKGs in every test case, indicating that our approach captures the specific differential epigenetic signals at cancer marker genes (Fig. 3a). Moreover, both for CMGs and HKGs, the Irene AUC values are higher than the AUC values computed using the dPC1+dPC2 of gene promoters only (Fig. 3a). The fact that the genes ranked higher in Irene suggests that a significant part of the altered epigenetic alteration arises from distal enhancer regions. We then validated these findings on the larger CMG and HKG gene sets, and we found the AUCs of CMGs are all significantly higher (one-tailed t-test p-value<0.01) than the AUCs of HKGs. (Fig. 3b,c)

**Figure 3:**
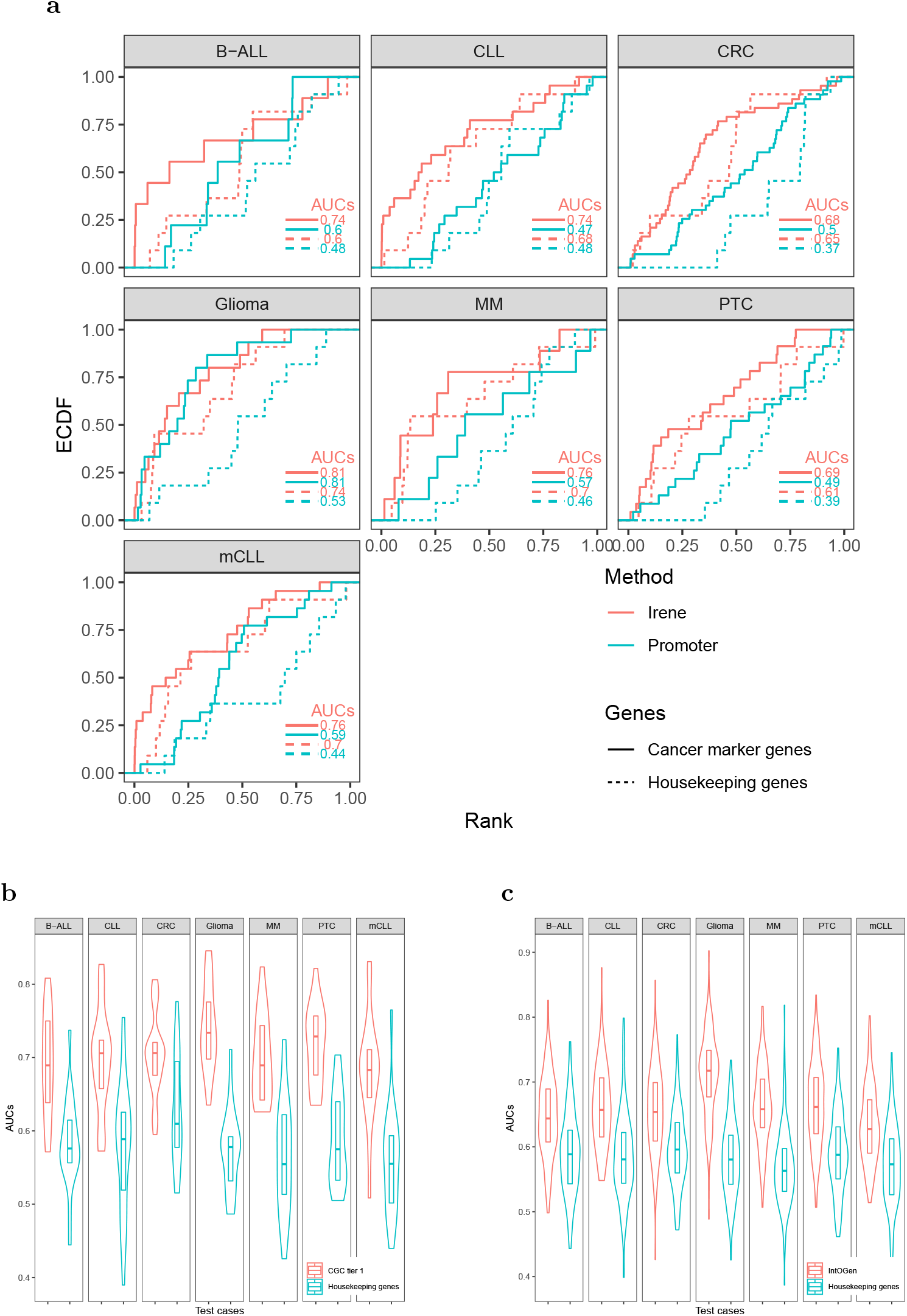
(a) ECDF curves regarding the cancer marker genes and housekeeping genes in seven test cases. The marker gene ranks using Irene scores (red) are compared against their ranks using the promoter scores (cyan). (b,c) Distribution of the AUC values using the CMG sets from Cancer Gene Census (Sondka et al., 2018) (b) and IntOGen (Gonzalez-Perez et al., 2013) (c) in the seven test cases, compared to randomly picked housekeeping genes to define equal size sets.

Some genes have a much high number of linked enhancers than others. To test whether this might bias the ranks of these genes, we performed 1000 degree-preserving random perturbations, which completely rewired the enhancer-promoter graph but maintaining the degree distribution. We used the high-confidence cancer marker genes in the benchmarking, and the AUCs with randomly assigned enhancers dropped 5-10% on average, indicating that the higher ranks of CMGs are not explained by their higher connectivity (Fig. 4)

**Figure 4:**
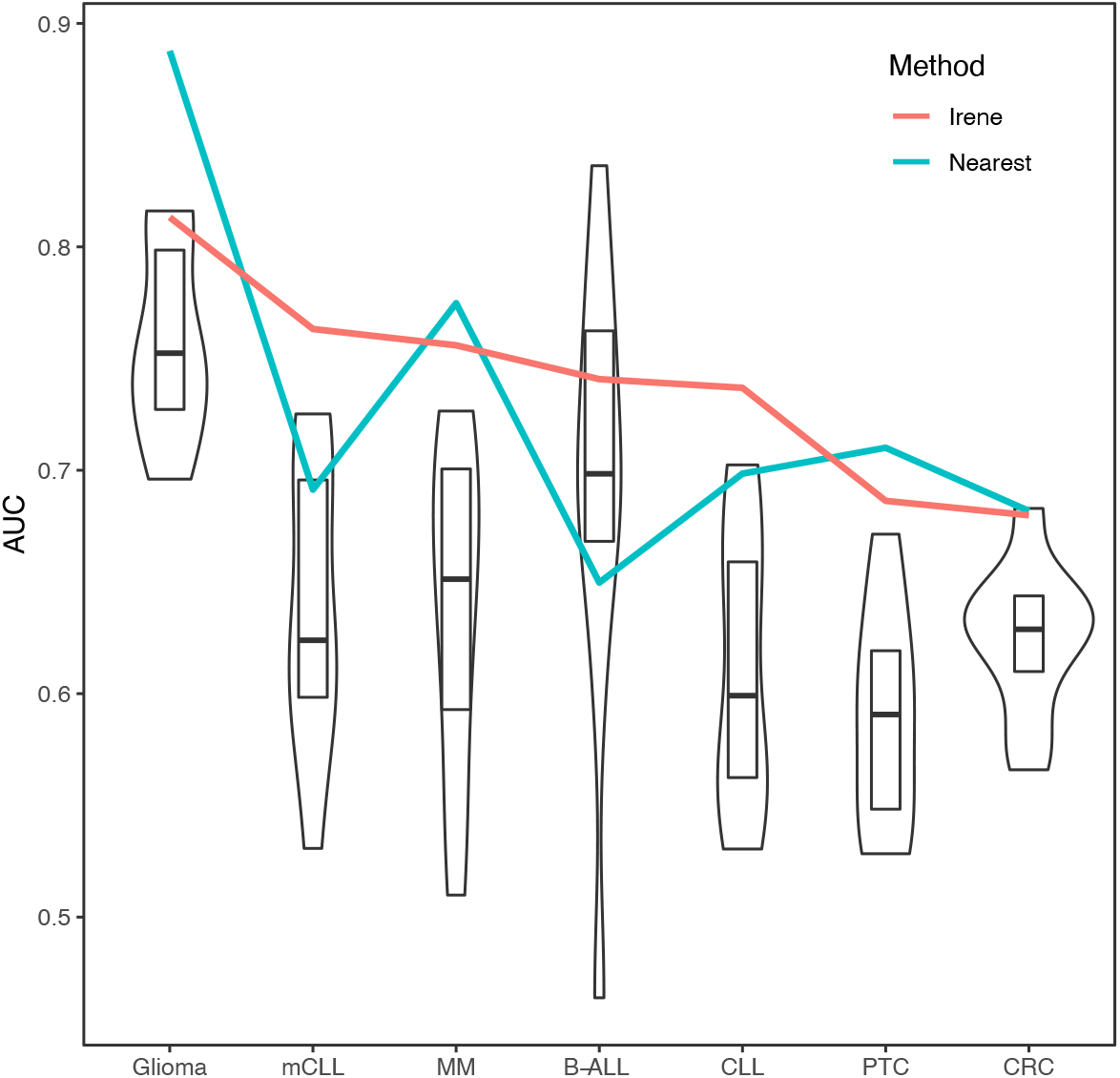
AUCs of ECDF curves of dPC1 ranks from randomized promoter-enhancer interactions. The boxplots indicate the 25%-75% quantile ranges from benchmarking each cancer marker gene set with 1000 different rewired promoter-enhancer networks, whereas the red lines show the AUCs with the original promoter-enhancer interactions from Irene using experimentally-detected interactions from 4DGenome (red), and interactions assigned using the nearest promoters (green).

Finally, we compared the target gene assignment provided by the 4DGenome database, which is based on experimental evidence, with the simpler nearest-gene assignment. As can be observed in Fig. 4, both approaches lead to comparable results, in line with recent reports indicating that the nearest gene assignment is reasonably effective in linking enhancers with target genes (Moore et al., 2020).

### Network analyses characterized the highly-ranked genes in the Irene and promoter list

We downloaded 184 KEGG pathways in KGML format and loaded them as directed graphs using KEGGgraph (Zhang and Wiemann, 2009). Then we took the top 15% genes from the Irene and promoter rank lists in each one of the seven test cases, and mapped the genes to the KEGG cancer signaling pathway (hsa05200). In total, the reference pathway contains 531 genes and 1,989 interactions, and on average 208 of the 531 genes are found in the Irene rank lists, while only 152 genes are found in the promoter rank lists. In addition, the Irene-ranked genes differ from promoter-ranked genes in both in-degrees and out-degrees of the nodes (Table 1). As the Irene nodes generally have higher in-degrees than out-degrees in the graph presentation of the reference pathway, implying the Irene genes are more often targeted by the other regulatory genes on their enhancers as they harbor more differential enhancers. We further examined the glioma signaling pathway (hsa05214) and found 19 genes from the Irene rank list and 10 genes from the promoter rank list in the glioma test case (Fig. 6). One common gene, *EGFR*, is in both lists and has been reported to undergo tight control through epigenetic regulation on both promoters and enhancers (Liu et al., 2015; McInerney et al., 2000; Jameson et al., 2019). Moreover, nine genes are present only in the Irene rank list, such as *CCND1*, which has been reported to be regulated by an estrogen-mediated enhancer (Eeckhoute et al., 2006). In conclusion, this analysis shows that the Irene methods provide a ranked gene list which is enriched for high-ranking, cancer-relevant genes.

**Figure 5:**
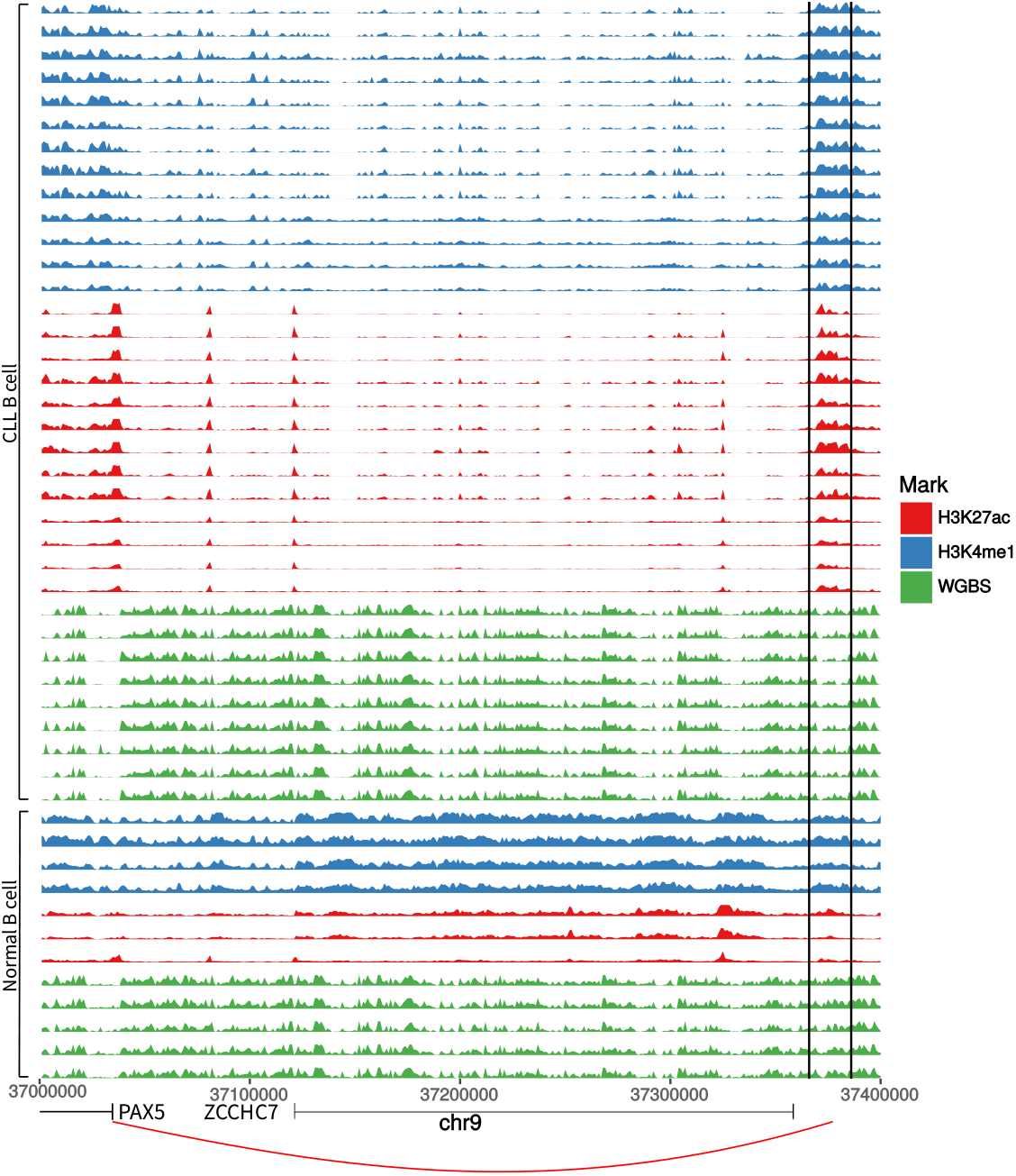
A known PAX5 enhancer in CLL exhibits hyperacetylation and hypomethylation. PAX5 enhancer positions in each track are surrounded by two black solid lines.

**Figure 6:**
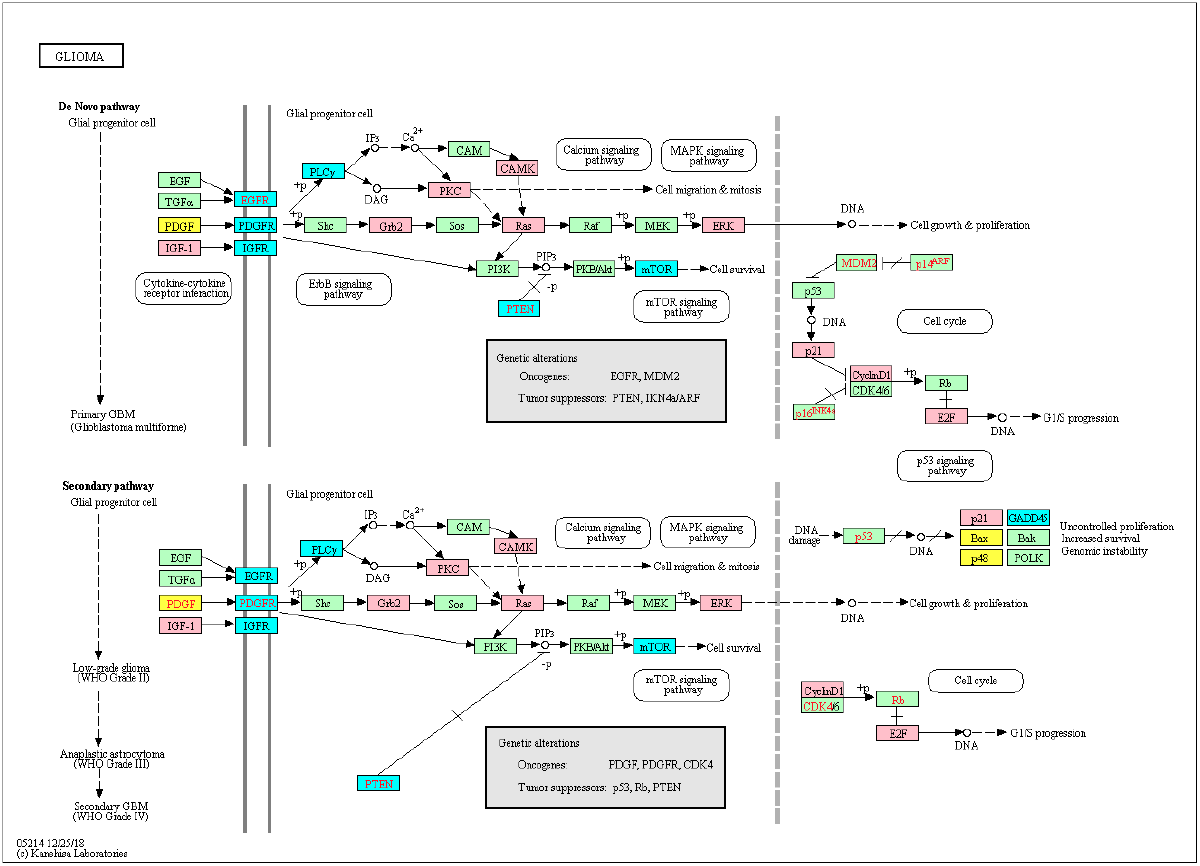
The top 25% genes from the Irene and promoter rank list are highlighted on the KEGG glioma signaling pathway. Pink: genes from the Irene list; Yellow: genes from the promoter list; Cyan: genes from both lists.

**Table 1:**
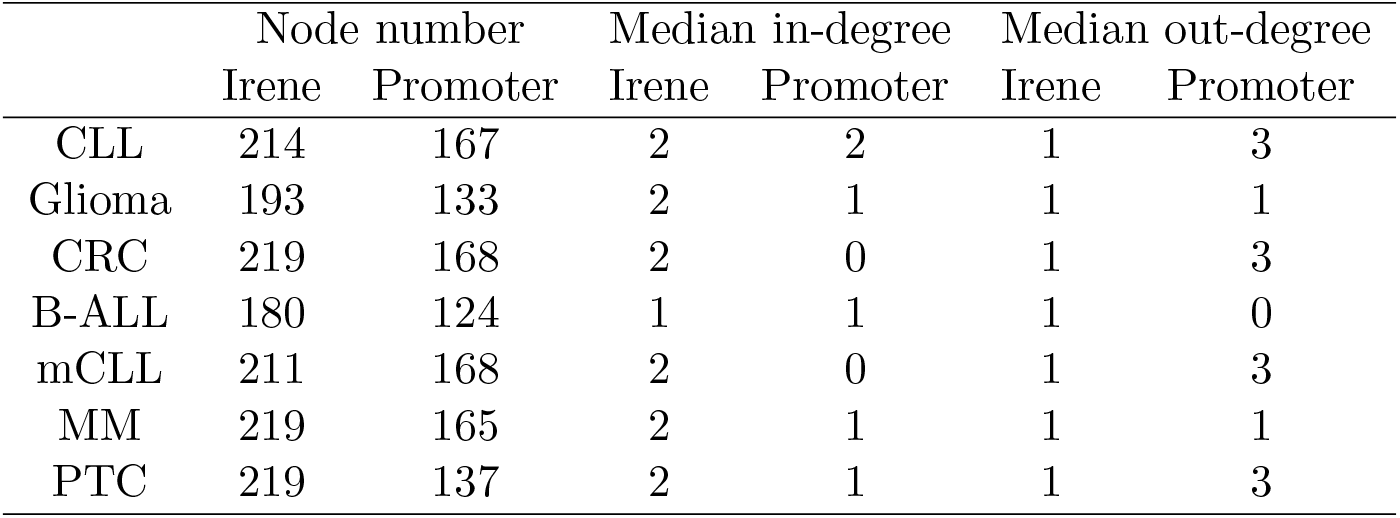
Graph properties in respect of the nodes from the Irene and promoter rank lists.

|  | Node number |  | Median in-degree |  | Median out-degree |  |
| --- | --- | --- | --- | --- | --- | --- |
|  | Irene | Promoter | Irene | Promoter | Irene | Promoter |
| CLL | 214 | 167 | 2 | 2 | 1 | 3 |
| Glioma | 193 | 133 | 2 | 1 | 1 | 1 |
| CRC | 219 | 168 | 2 | 0 | 1 | 3 |
| B-ALL | 180 | 124 | 1 | 1 | 1 | 0 |
| mCLL | 211 | 168 | 2 | 0 | 1 | 3 |
| MM | 219 | 165 | 2 | 1 | 1 | 1 |
| PTC | 219 | 137 | 2 | 1 | 1 | 3 |

## Discussion

From the above benchmarking on seven cancer test case studies, we showed that Irene is a more comprehensive approach comparing to the current frequently used approaches such as separate ranking gene promoters and enhancers. This highlights the importance of epigenetic regulation through distant enhancer regions. Using Irene, users can not only discover the genes which show significantly epigenetic alterations on their promoters, but also the ones which are connected with strong epigenetic modifications on distal interacting enhancers, which facilitates the discovery of potential epigenetic marker genes. On the other hand, by interpreting the higher-ranked genes mapped to the existing pathways, the user may also find the enhancers of interests from their differential epigenetic modifications. For example, we found the *PAX5* gene to have a significantly higher rank in the Irene list compared to the promoter-only list in the two CLL case studies, which implies that *PAX5* is extensively regulated by enhancers. *PAX5* is a key TF involved in B-cell development, and its promoters have no significant epigenetic alterations in the CLL case studies. However, this gene is associated with several hyperacetylated and hypomethylated distal enhancers, one of which is located at 330 kilobases (kb) upstream of the PAX5 TSS, and has been also found as extensively mutated in CLL (Puente et al., 2015) (Fig. 5). The deletion of this enhancer resulted in a 40% reduction in the expression of *PAX5* expression and chromatin interaction of this enhancer and *PAX5* has been proven from chromosome conformation capture sequencing (4C-Seq) analysis (Puente et al., 2015). The main difficulty of this study is obtaining cell-type specific enhancer-promoter interactions, as the high-resolution chromatin interaction map for the cancer cells is currently not available. We have tested two alternative approaches in this study, using either the experimentally validated chromatin interaction or distance-based interactions. The performance of the above two approaches are similar (Fig. 4). We believe better performance can be achieved when cell-type specific enhancer-promoter interactions are available in the future, and using Irene, user can replace the interaction map with a more specific one when applicable.

## Conclusions

Genome-wide integrative epigenetic analysis is challenging and essential in many comparative studies. As far as we know, Irene is the first tool that integrates quantitive and genome context information in the differential epigenetic analysis. Applying this tool to well-characterized test cases, it detects a number of candidate genes with significant epigenetic alterations, and comprehensive benchmarking validated these findings in cancer studies. As the accumulation of the epigenomic datasets, the computational approaches employed in this study would be highly relevant in both comparative and integrative analysis of the epigenetic landscape. The discovery of novel epigenetic targets in cancers, not only unfolds the fundamental mechanisms in tumorigenesis and development, but also serves as an emerging resource for molecular diagnosis and treatment.

## Material and methods

### Data Preparation

#### Retrieving Epigenetic Modification And Chromatin Interaction Datasets

Genome-wide ChIP-seq data are downloaded in BigWig format from NIH Roadmap Epigenomics (Bernstein et al., 2010), Blueprint (Adams et al., 2012), and the International Human Epigenome Consortium (IHEC) (Stunnenberg et al., 2016). We selected the six most frequently studied histone marks: H3K27ac, H3K27me3, H3K36me3, H3K4me1, H3K4me3, and H3K9me3. These resources allow us to investigate the histone modification differences between tumor and normal tissues (Supplementary Table 1). For restricting the comparisons to the genomic loci of interests (promoters and enhancers), we downloaded the GRCh37 and GRCh38 coordinates of promoters from the eukaryotic promoter database (EPD) (Dreos et al., 2013), and the promoter interacting regions (PIRs) from the 4DGenome database (Teng et al., 2015). We treated the PIRs as potential enhancer regions, and filtered for tissue-specific enhancers by requiring the presence of H3K4me1 or H3K27ac peaks (peak calls provided in the Supplementary Table 1) in at least two samples from either tumor or normal tissues. By doing this, we enrich for cell-type specific PIRs, which show a tissue-driven clustering (Supplementary Fig. 1). The promoter coordinates were extended to ±1000 base pairs around the original coordinates. The sum of the numeric values from the BigWig blocks which overlap with the promoter and interacting regions are available from our project homepage. To build the relationships between and enhancers and promoters, we also download all the experimentally validated chromatin interaction datasets in various human tissues from 4DGenome.

#### Defining disease and control datasets

We used histone modification datasets from seven cancer types in this study, i.e., B-ALL, CRC, glioma, MM, PTC, CLL, and mCLL from the Blueprint and IHEC consortia. For each cancer dataset, we paired it with the available dataset from the healthy tissue which the cancer is most likely originated from. For example, the B-ALL, CLL, MM were all compared against the healthy B-cells in our design (see Supplementary Table 1 for the pairs of normal/tumors used).

#### Definition of cancer marker genes and housekeeping genes

We evaluated our algorithm on a small set of high-confidence cancer marker genes (CMGs), which is based on the tier-1 genes of the corresponding tissues from the Cancer Gene Consensus (CGC-t1) (Sondka et al., 2018) (Supplementary Table 2). As a negative control, we retrieved 11 housekeeping genes (HKG) that show constant expression profiles in RNA-Seq studies (Eisenberg and Levanon, 2013). To validate our findings on independent, larger datasets of CMGs and HKGs, we compiled two additional CMG lists containing 303 CMGs (11-63 for each cancer type) of 21 cancer types from the Cancer Gene Consensus (Sondka et al., 2018), and 546 CMGs (11-55 each cancer type) of 129 cancer types from IntOGen (Gonzalez-Perez et al., 2013). We picked random control genes of equal sizes as the CMGs among 515 housekeeping genes of four different categories (RNA polymerase, ATPase, NADH dehydrogenase, SLC transporters).

### Data processing procedures

#### Combining histone marks

The epigenetic intensities on regulatory elements were summarized on a 1-kilobase-scale, and then quantile normalized. For using the differential principal component analyses (dPCA), the data were transformed into normal distributions using Box-Cox transformation before quantile normalization. We implemented an R wrapper function for dPCA in our tool, which takes the mean differences of the normalized ChIP-Seq signals in each genomic locus between two biological conditions as input, and returns the differential principal components (dPCs) from dPCA. The definition of dPCs varies between the test cases (Fig. 2a). The largest variances of the positive and negative histone mark components are captured by dPC1 and dPC2 in our test case studies (Fig. 2b). Therefore we selected the sum of the absolute values of the first two dPCs for representing the overall differences of these epigenetic marks.

#### Promoter-enhancer interaction analyses

In our approach, the enhancer-promoter relationships are described as a weighted bipartite graph, in which both enhancers and promoters are represented as vertices, and edges are directed from enhancers to their target promoters (Fig. 1 (Step 1)). The weights of the vertices are defined as the sum of the absolute values of the first two dPCs when combining multiple epigenetic marks, or the absolute value of the difference if a single epigenetic mark is considered. We adopt an algorithm called “PageRank”, which is originally designed for evaluating the importance of web pages (Brin and Page, 1998), for ranking the magnitude of epigenetic alterations in each gene. We use the “personalized” PageRank implemented in igraph (Rye et al., 2011) to summarize the weights of one promoter and its connected enhancers into a unique meta-gene score (Fig. 1 (Step 2)). Since our enhancer-promoter network is a directed graph, all the enhancer weights will eventually be attributed to their target promoter using PageRank, yielding a unified score for each gene, which can be used to rank the genes. Overall, there are ~ 251, 000 promoter interacting fragments in the promoter-enhancer interaction networks in our case studies, which is 8.5 times the number of promoters in the networks. The number of the interacting fragments targeting a gene varies from none to 227, and on average, 21 interacting fragments are targeting a promoter in the networks.

#### Scoring ranked lists

Using the gene ranks computed as described in the previous section, we can now evaluate the enrichment of a specific gene set 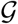 in the ranked list by computing the empirical cumulative distribution function (ECDF) obtained ranking the genes in decreasing order based on the previously described rank, and summing the indicator function

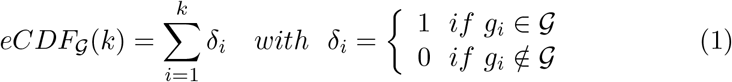

We use the area under the curve (AUC) as a measure of the enrichment of the gene set 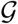, with AUC=0.5 corresponding to a random distribution of the genes in 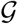 inside the ranked list.

## Data Deposition

Results for the test cases can be found under https://uni-hd.github.io/Irene/. The R package is available at https://github.com/uni-hd/ Irene. We also designed a web interface that allows users to trace back the epigenetic alterations of every enhancer and promoter, as well as every sample which is used for computing the score. We use Rmarkdown to generate static HTML pages and created a web site for presenting the results from our test case studies, which can also be found under the project home page. Users may also take advantage of this function to create a report that highlights a few genes of their interests and share the studies with the audience.

## Author’s contributions

CH designed and supervised this project. QW and CH drafted the manuscript. QW wrote and tested the software. YW and TV participated in software testing. YW, TV, and RE revised the manuscript.

## Acknowledgments

This work was supported by the BMBF-funded PRECISe project (#031L0076A).

