## Supplementary Figure 1 for "Integrative Ranking Of Enhancer Networks Facilitates The Discovery Of Epigenetic Markers In Cancer"

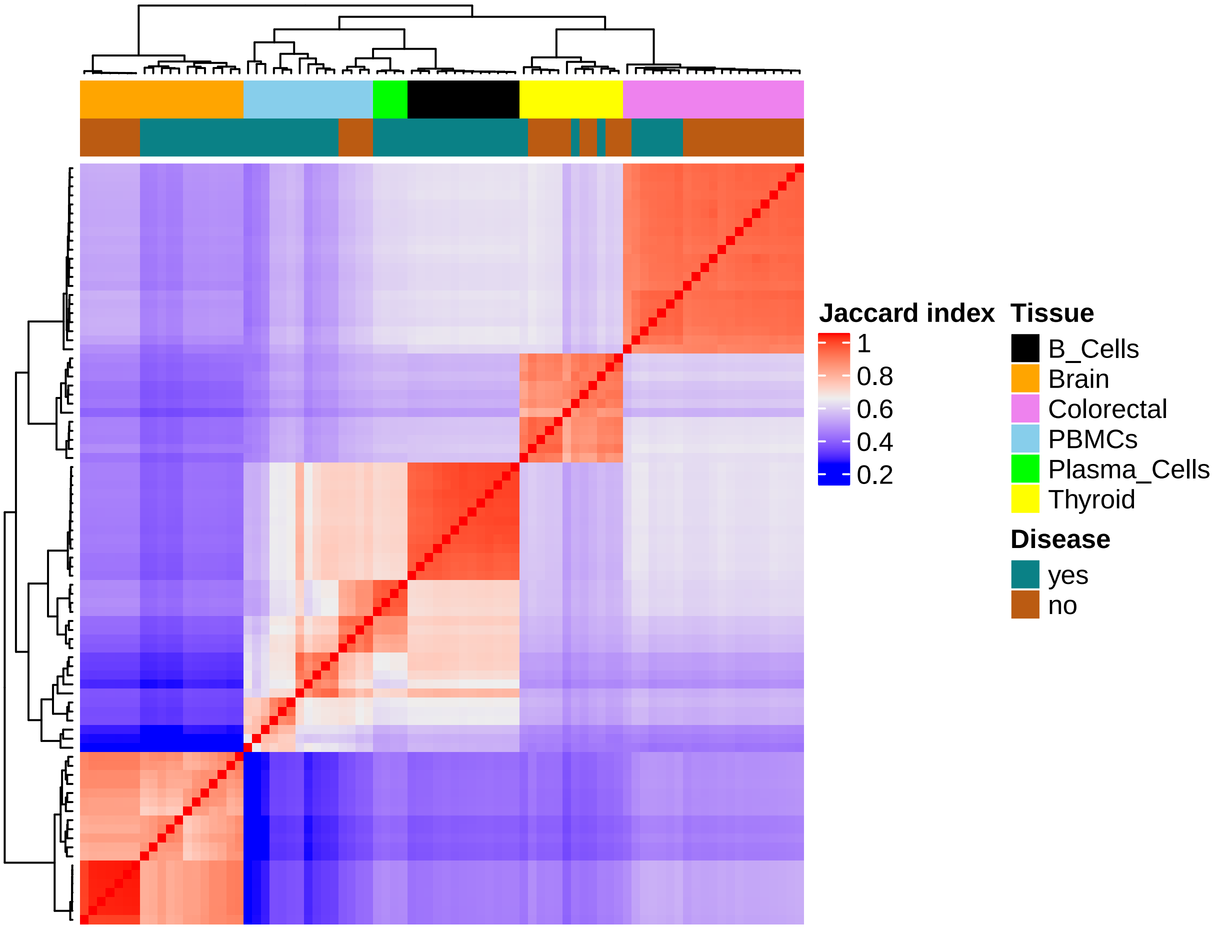


Supplementary Figure 1: **Clustering of common PIRs between samples shows enrichment of cell-type specificity.** We intersect the peaks obtained from enhancers marks (H3K4me1 and/or H3K27ac) in each sample with the list of PIRs (see "Data preparation" section), and computed the Jaccard index using the intersection and union of filtered PIRs between samples.
